# The effects of a 10-day partial sleep deprivation and the following recovery on cognitive functioning – a behavioural and EEG study

**DOI:** 10.1101/666396

**Authors:** Anna M. Beres, Aleksandra Domagalik, Jeremi K. Ochab, Katarzyna Oleś, Halszka Ogińska, Magdalena Fąfrowicz, Tadeusz Marek, Ewa Gudowska-Nowak, Maciej A. Nowak, Dante Chialvo, Jerzy Szwed

## Abstract

Sleep deprivation is an important societal problem that affects millions of people around the world on a daily basis. Our study aimed to examine the impact of a partial sleep restriction and following recovery processes on cognitive information processing, as evaluated by the Stroop test. We tested 15 participants over a period of 21 consecutive days, divided into 3 sleep conditions: 4 days of typical daily routine (baseline, ‘base’), 10 days of partial sleep deprivation (‘SD’), and 7 days of recovery (‘rec’). Each day, participants took part in an EEG experiment in which they performed a Stroop test, lasting for about 30 minutes, that required them to make an appropriate response to congruent and incongruent stimuli. Additionally, every day they answered a number of questions regarding their subjective levels of sleepiness and mood. During the whole period of 21 days, participants’ spontaneous locomotor activity was measured with the use of actigraphy. We have found behavioural and neural changes associated with different sleep conditions, such that the 10-day period of partial sleep restriction was linked with poorer behavioural performance on the Stroop test and an attenuated P300 neural response, compared to the baseline, followed by the observation of slow and gradual return in the period of recovery. This study, the first longitudinal study of its kind, shows that partial sleep deprivation has detrimental, long-term consequences on both behavioural and neural levels. This adds to the growing body of literature on this important issue in modern societies.

**Summary:** Sleep deprivation, a world-wide problem in the 21st century, is associated with a number of complications, such as motor vehicle accidents (Lyznicki et al., 1998; Goel et al., 2009), medical errors (Barger et al., 2006), poorer health (in Colten & Altevogt, 2006), as well as cognitive deficits including problems with working memory and attention (van Dongen et al., 2003; Lim & Dinges, 2008). While total sleep deprivation (that is, a complete lack of sleep in a 24-hour period) is usually reserved only to certain professions (such as medical doctors), partial sleep restriction (that is, reducing one’s sleep time in a 24-hour period to fewer hours than typically required) is world-wide and affects a large proportion of the population across the globe. Taking this global impact into account, and thus increasing our understanding of the neurophysiological and cognitive processes that are linked with partial sleep deprivation, could largely inform the public discussion over what kind of impact, if any, restricting our sleep has on our daily functioning. This 21-day long EEG study investigated the effects of a prolonged (10-day) sleep restriction, and the recovery processes (over a 7-day period) that followed. Each day participants performed a Stroop test, known to measure attentional levels, and completed a number of sleep-related questionnaires. We have found that while behavioural responses are easier to recover, the neurophysiological responses are heavily affected after a period of sleep deprivation, with one week of recovery being insufficient to return to a pre-testing performance of an individual.

## Introduction

Sleep is a crucial aspect of human’s mental and physical well-being. Negative cognitive consequences linked with sleep deprivation present worldwide, and include sensory perception, emotion regulation, attention, executive functioning, and memory (Goel et al., 2013; Landoldt et al., 2014; Lowe et al., 2017; Medic et al., 2017). Even though in modern societies partial sleep restriction (when an individual often sleeps less than required for their optimal functioning, Cirelli et al., 2016) seems to be a growing problem, it is a less studied phenomenon than total (also known as acute, Cirelli et al., 2016) sleep deprivation (TSD, a complete lack of sleep for 1-2 nights) (Banks & Dinges, 2007). Some research suggests that both forms of sleep loss may have similar consequences (e.g. Banks & Dinges, 2007; van Dongen et al., 2003). Van Dongen et al. (2003) showed that chronic restriction of sleep to six hours or less per night over 14 consecutive nights results in impaired cognitive performance, including lapses in behavioural alertness and working memory performance, equivalent to up to two nights of TSD. Unlike cognitive performance measures, the subjective sleepiness response to sleep loss over days was different for partial and total sleep deprivation. The study by Sallinen and colleagues (2013), which investigated chronic partial sleep deprivation (5 hours of sleep for 5 consecutive nights), showed that self-perceived task performance, sleepiness and mental fatigue were all impaired during the sleep restriction phase and returned to baseline in the recovery phase, while at the same time self-perceived mental effort, tension, task difficulty and pace were not affected by the sleep loss. The effects of sleep loss can also be observed at the neural level (e.g., Belenky et al., 2003; Hsieh et al., 2010; Lo et al., 2012; Renn & Cote, 2013; van Dongen et al., 2003), with changes in the amplitude and latencies observed in a P300 component of the ERPs, known to index information processing, such as the individual’s reaction to novelty and improbability of stimuli in attention tasks with manual reactions.

However, very few studies have focused on the actual mechanisms involved in the recovery processes following reduced sleep. The majority of those that have, looked either at total sleep deprivation (Lamond et al., 2007) or included only a short time of recovery (Belenky et al., 2003; Caldwell & Caldwell, 1997; Dinges et al., 1997; van Dongen et al., 2003), usually between 1-3 days. Therefore, the purpose of the present study was to examine the behavioural and neural consequences of chronic partial sleep deprivation in healthy individuals, and subsequently their ability to recover following a prolonged period of controlled sleep restriction.

We designed a study that included periods of 4 days of normal sleep, 10 days of sleep deprivation and one week of recovery. We applied a Stroop paradigm and EEG recording in order to investigate higher cognitive functioning at behavioural and neural levels. Actigraphy was used for control of sleep-wake cycle. Additionally, subjective scales were used to asses sleepiness level and mood (positive and negative).

## Methods

### Participants

15 healthy adults (14 females, age 21.47±1.36 y.o.) participated in the study. All participants were healthy and declared that typically they sleep no less than 7 hours a night. The study was approved by the Bioethics Committee at the Jagiellonian University and all participants gave their written consent. They were financially reimbursed for their time.

### Procedure

The whole study lasted for 21 consecutive days. The sessions were divided into three sleep conditions: a 4-day period of unrestricted sleep (baseline, ‘base’), a 10-day period of daily sleep curtailment by ~30% of individual baseline sleep length (sleep deprivation. ‘SD’), followed by a 7-day period of recovery (‘rec’) (See Figure 1, below, for a graphical representation of the study design). Each day, participants’ subjective sleepiness levels were assessed with the use of the Karolinska Sleepiness Scale (KSS) and their mood was assessed with the use of Positive and Negative Affect Schedule (PANAS) questionnaire. Every day they also performed a Stroop test while their brain activity was being monitored with an EEG (64 channels, Geodesic EGI System 300, USA). Each participant had undergone the testing procedures at the same time every day. Additionally, for the entire 21 days of the study, participants’ sleep timing and duration was controlled with actigraphy (Micro Motionlogger Sleep Watch, Ambulatory Monitoring, Inc., Ardsley, NY).

**Fig. 1.**
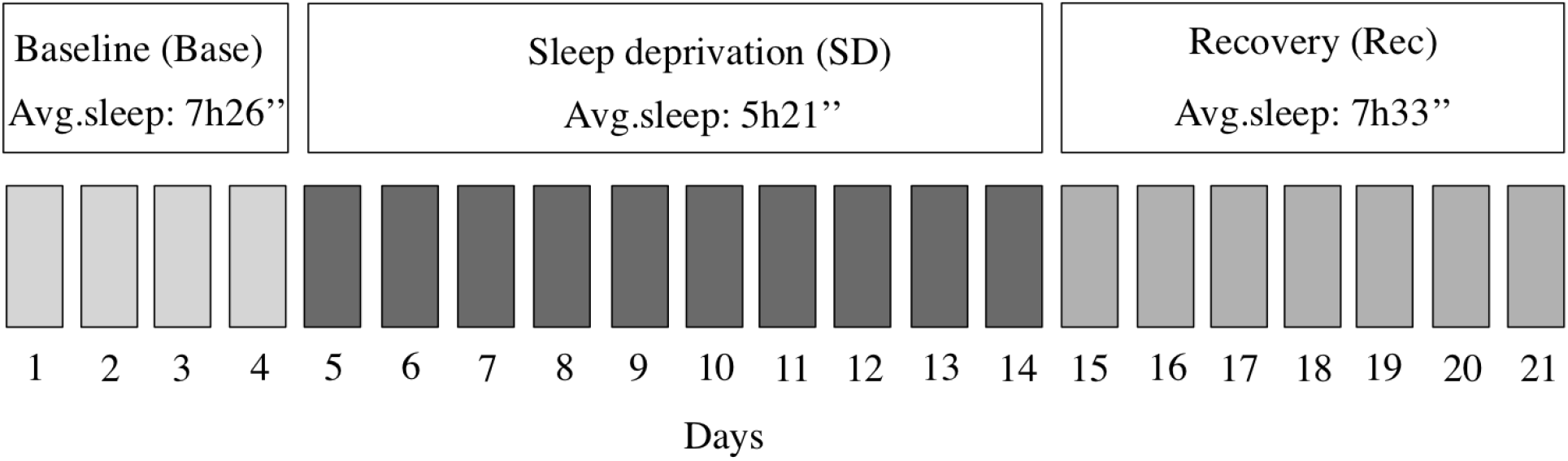
Study experimental design, showing number of days in each of the experimental sleep conditions, and the average sleep duration in each of the 3 conditions.

### Task

Each day, participants performed a Stroop test, with 4 different colours (red, blue, green, and yellow). There were a total of 432 randomly shuffled stimuli (half of them congruent), presented in 3 separate blocks with a short break in between each block. The task was to indicate with a corresponding button on a keyboard whether the presented stimulus is congruent (that is, whether the name of the colour and ink colour are the same) or incongruent (where the name and the ink of the colour are different). The inter-stimulus interval between each stimulus was between 1500 ms and 3500 ms (in steps of 400 ms), with 2500 ms on average. The entire task duration was 22 minutes and 11 seconds (22.19 minutes on average). Participants took part in one training session before beginning the experiment. See Fig. 2 for an example of congruent and incongruent trials in the Stroop task.

**Fig. 2.**
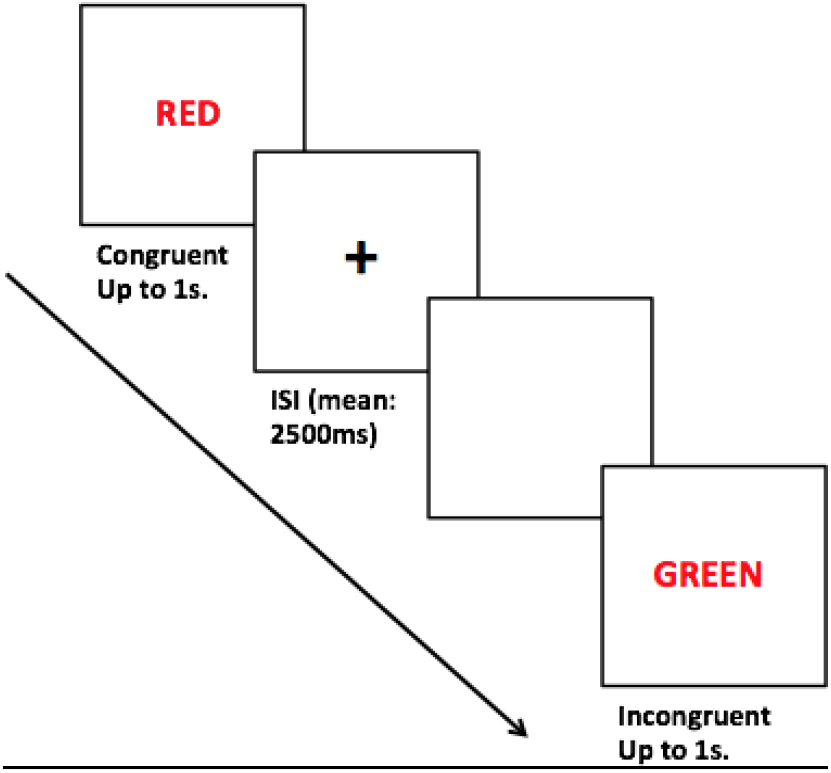
A graphic representation of a Stroop task.

### Subjective measures

Karolinska Sleepiness Scale (KSS – Åkerstedt & Gillberg, 1990) is a classic tool used in self-assessments of actual sleepiness level, with good psychometric characteristics and an extensive body of literature linking its scores to objective measures. This simple scale is scored from 1 (extremely alert) to 9 (extremely sleepy, fighting with sleep). The Positive and Negative Affect Schedule (PANAS – Watson et al., 1988) is a self-report measure of mood comprising two sub-scales, each containing ten items. Positive affect (PA) reflects the extent to which a person feels enthusiastic, active and alert. High scores indicate a high level of concentration, energy and commitment. Negative affect (NA) is a general dimension of subjective distress covering a variety of aversive mood states – anger, disgust, fear, nervousness etc. The items are scored from 1 (‘very slightly or not at all’) to 5 (‘extremely’).

### EEG recording and analysis

Data were digitally filtered (0.5–35 Hz), average references were computed, bad channels and time epochs with artefacts removed. An Independent Component Analysis (ICA) was used to separate and remove physiological artefacts, including saccade-related spike potentials (Delorme and Makeig, 2004). Event-related potentials (ERPs), locked to the target onset, were obtained for congruent and incongruent trials from the Cz recording site. Data was not analysed on a day-by-day basis, but rather averaged across the particular sleep conditions of the experiment for each subject, such that the first 4 days resulted in ‘baseline’, the following 10 days in ‘sleep deprivation’, and the final 7 days in ‘recovery’ periods.

For the ERP measures, segments were created from 200 ms before to 1000 ms after the onset of the target stimulus. The trials associated with each block were averaged for each participant. Afterwards, the data was subjected to a 3 × 2 ANOVA analyses, with ‘sleep condition’ (baseline, SD, and recovery) and ‘congruency’ (congruent, incongruent) as factors.

As expected in a Stroop experiment, with the focus on the P300 effects, the strongest differences occurred within the centro-parietal sites. For the ERP analyses, time window of 400-600 ms was chosen for the analyses, with the strongest effects present at the E34 site (centroparietal region).

All statistical tests were performed using SPSS software (SPSS Corp., released 2016. IBM SPSS Statistics for Macintosh, version 24.0. Armonk, NY:IBM Corp.)

## Results

### Participant’s sleep

On average, participants slept 7 hours and 26 minutes (± 1h 7”) during the baseline, which then reduced by about 28% in the SD period, to 5 hours and 21 minutes (± 33”). In the recovery period, participants’ sleep duration increased to 7 hours and 33 minutes (± 34”) on average. There was a statistically significant decrease in sleep duration in the SD period, compared to the baseline (p < 0.001) and a significant increase in the recovery period compared to SD (p < 0.001). There was no significant difference in sleep duration between baseline and recovery.

### Handling of the data

One participant had to be removed from the behavioural analyses (n = 14) and four from the EEG analyses (n = 11) due to insufficient quality of the data.

Repeated measures ANOVA, with two factors: ‘sleep condition’ (3 levels: baseline, sleep deprivation, recovery) and ‘congruency’ (2 levels: congruent, incongruent) was performed. Below we present the results of the subjective measures, Stroop task performance and EEG data.

All statistically significant results have been adjusted for multiple comparisons using Bonferroni computations. All effects are reported as significant at p < 0.05.

### Subjective measures

Reported sleepiness was significantly higher in the period of sleep deprivation, compared to both the baseline and recovery periods, F(2,13) = 49.45, p < 0.001, partial η2 = 0.884. (Fig. 3)

**Figure 3.**
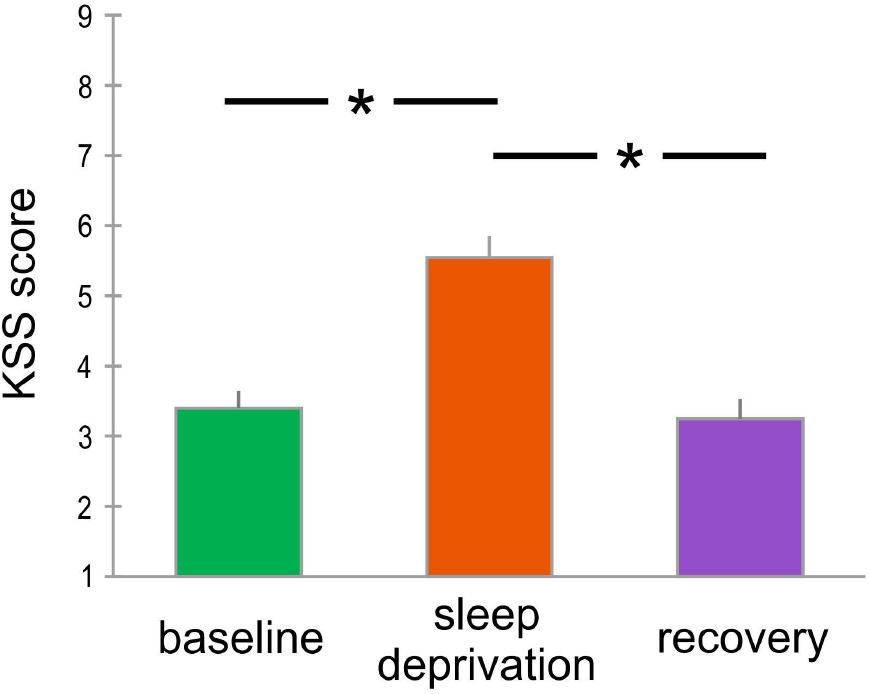
Reported subjective levels of sleepiness in each of the experimental conditions (n = 14).

Participants reported significantly higher negative affect in the sleep deprivation period, compared to the other two phases, F(2,13) = 4.519, p < 0.032, partial η2 = 0.365. This reflects a difference between baseline and SD (p = 0.02) and SD and recovery (p = 0.04) phases. (Fig. 4)

**Figure 4.**
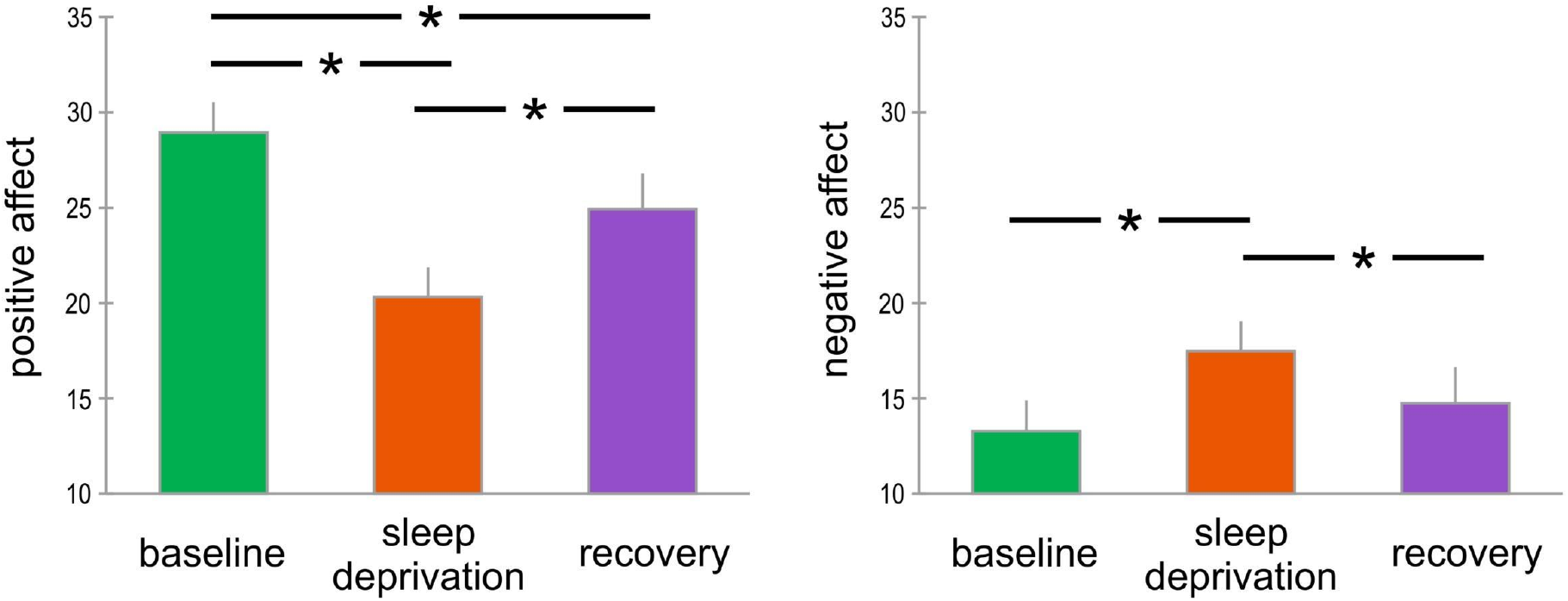
Reported subjective positive and negative mood in each of the three experimental conditions (n = 14).

Subjects also reported lower positive affect in the sleep deprivation period, compared to the other two phases, F(2,13) = 27.7, p < 0.001, partial η2 = 0.81. This reflected a difference not only between baseline and SD (p < 0.001) and SD and recovery (p = 0.006) conditions, but also, interestingly, between the baseline and the recovery phase (p = 0.011), meaning that despite some increase in the reported feelings of positivity in the period of recovery, those levels were still significantly lower than at the beginning of the study. (Fig. 4)

**Figure 4.**
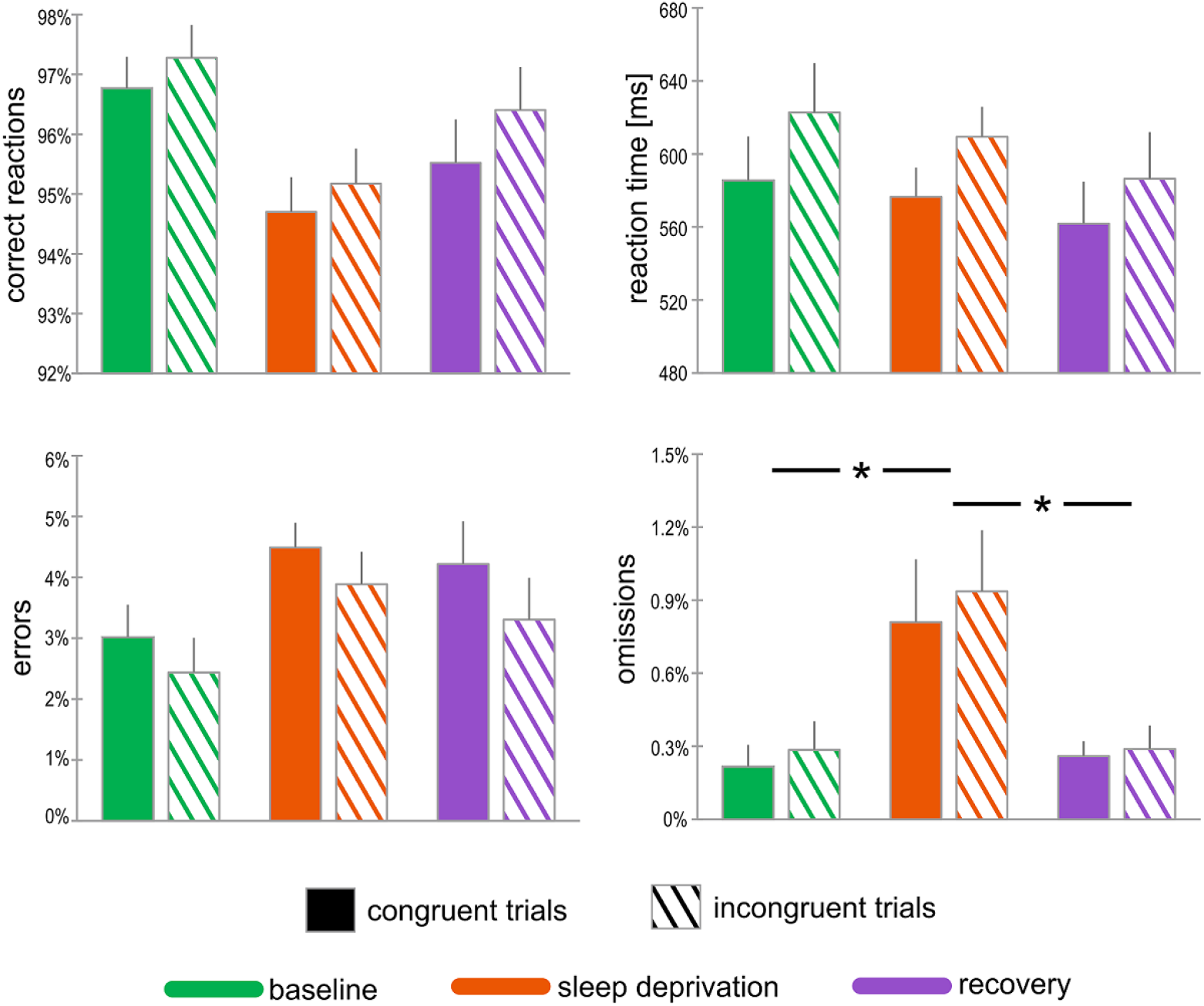
Behavioural results (n = 14) showing the differences between the three sleep conditions (baseline, SD, recovery) for *correct responses* and *reaction times to correct responses* (top row) as well as the number of *errors* and *omissions* (bottom row). Bars denote means, and whiskers denote standard errors.

### Behavioural results

Table 1, below, lists participant’s correct reactions, reaction times (RTs), errors and omissions to congruent and incongruent stimuli in each of the 3 experimental conditions. Figure 4 (below) shows a graphical representation of those results.

**Table 1.**
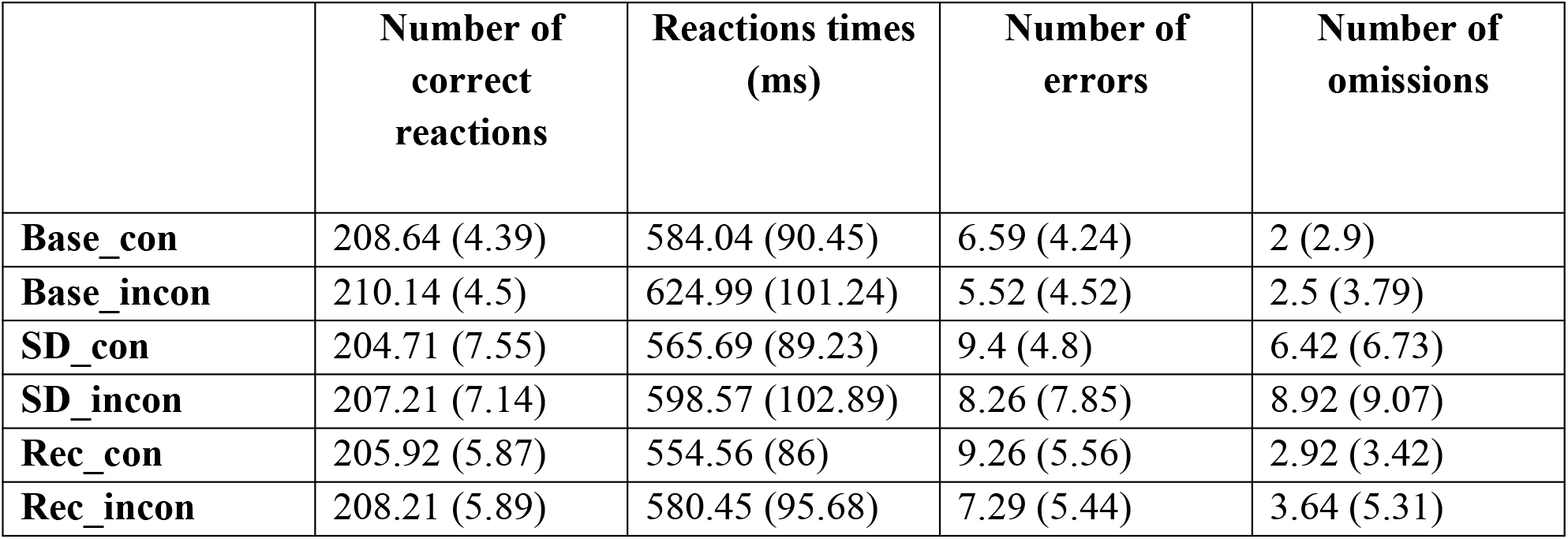
Mean and standard deviation (in brackets) values for congruent (_con) and incongruent (_incon) stimuli in the 3 different sleep conditions (n=14).

|  | **Number of**<br>**correct**<br>**reactions** | **Reactions times**<br>**(ms)** | **Number of**<br>**errors** | **Number of**<br>**omissions** |
| --- | --- | --- | --- | --- |
| **Base_con** | 208.64 (4.39) | 584.04 (90.45) | 6.59 (4.24) | 2 (2.9) |
| **Base_incon** | 210.14 (4.5) | 624.99 (101.24) | 5.52 (4.52) | 2.5 (3.79) |
| **SD_con** | 204.71 (7.55) | 565.69 (89.23) | 9.4 (4.8) | 6.42 (6.73) |
| **SD_incon** | 207.21 (7.14) | 598.57 (102.89) | 8.26 (7.85) | 8.92 (9.07) |
| **Rec_con** | 205.92 (5.87) | 554.56 (86) | 9.26 (5.56) | 2.92 (3.42) |
| **Rec_incon** | 208.21 (5.89) | 580.45 (95.68) | 7.29 (5.44) | 3.64 (5.31) |

#### Correct reactions

Although there was a trend for the participants to provide fewer correct responses in the SD period compared to the baseline, and for those to slightly increase again in the recovery period, this difference failed to reach statistical significance (F(2,13) = 2.938, p = 0.07, partial η2 = 0.184). There was a main effect of ‘congruency’ (F(1,13) = 5.786, p = 0.03, partial η2 = 0.308), such that participants provided more correct responses to incongruent, rather than congruent stimuli. No interaction was found between the two main factors (F(2,13) = 0.544, p = 0.58, partial η2 = 0.040).

#### Correct reactions – reaction times

There was no significant main effect of ‘sleep condition’, (F(2,13) = 2.995, p = 0.06, partial η2 = 0.187). There was a main effect of ‘congruency’ (F(1,13) = 33.294, p < 0.001, partial η2 = 0.719), such that participants responded quicker to congruent, as opposed to incongruent stimuli as well as an interaction between ‘sleep condition’ and ‘congruency’ (F(2,13) = 4.749, p = 0.017, partial η2 = 0.268), such that participants’ reaction times to both congruent as well as incongruent stimuli decreased with each subsequent sleep condition, compared to the baseline.

#### Errors

There were no statistically significant main effects nor interactions in terms of a number of incorrect responses (all p values > 0.05).

#### Omissions

We have found a significant main effect of ‘sleep condition’, F(2,13) = 7.252, p = 0.003, partial η2 = 0.358, which reflects the difference between levels 1 and 2 (that is, baseline and SD) with p = 0.03 and levels 2 and 3 (that is, SD and recovery) with p = 0.04, such that there were significantly more omissions (irrespective of congruency) in the period of sleep deprivation, compared to baseline and to recovery periods. There was no significant main effect of ‘congruency’ (F(1,13) = 4.057, p = 0.065, partial η2 = 0.238), which shows that there was no difference in the omissions rates of congruent and incongruent stimuli. There was no significant interaction between the ‘sleep condition’ and ‘congruency’ factors, F(2,14) = 0.105, p = 0.06, partial η2 = 0.193.

### EEG results

There was a significant main effect of the ‘sleep condition’, F(2,10) = 4.878, p=0.01, partial η2 = 0.328, and of ‘congruency’, F(1,10) = 15.612, p = 0.00, partial η2 = 0.610. There was no significant interaction between the two factors, F(2,10) = 0.105, p = 0.900, partial η2 = 0.010.

The significant main effect of the ‘sleep condition’ reflects a significant difference between levels 1 and 2 (that is, baseline and sleep deprivation, p = 0.02) and 1 and 3 (that is, baseline and recovery, p = 0.04). There was no significant difference between levels 2 and 3 (that is, sleep deprivation and recovery, p = 0.80). See Figure 5 for the ERP waveforms elicited in response to congruent (A) and incongruent (B) stimuli.

**Figure 5.**
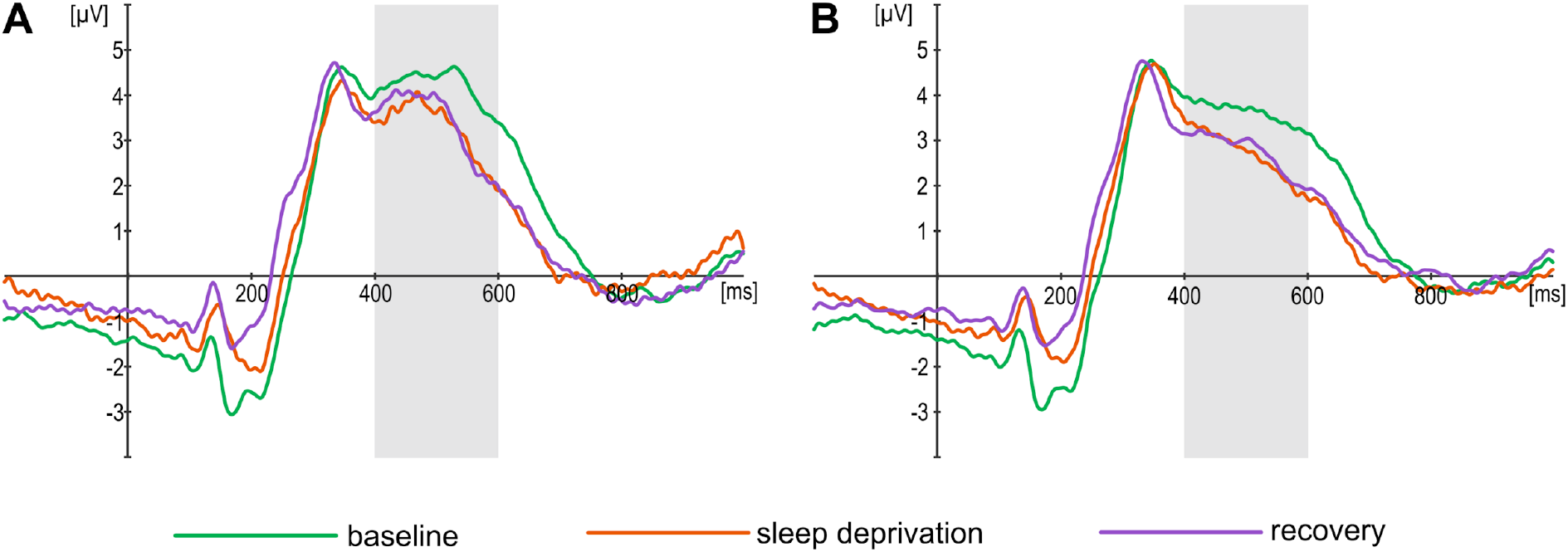
ERPs to congruent (A) and incongruent (B) stimuli in the Stroop test (n=11).

Subjective measures and behavioural data for each day of the experiment are presented in supplementary material.

## Discussion

The aim of this study was to investigate the differences in participants’ performance on a Stroop task while undergoing a longitudinal study of sleep restriction. In particular, we were interested in observing the possible neural and behavioural differences between the three sleep conditions of the experiment: baseline, sleep deprivation, and recovery. Finally, we were most interested in the processes involved in the recovery period, that is answering the question whether a period of 7 days of unrestricted sleep would be sufficient in regaining neural and behavioural patterns as seen in the baseline.

### Behavioural effects

As expected, the distributions of reaction times for congruent stimuli show smaller means compared to incongruent ones, in all three sleep conditions. What is surprising however, is that the accuracy of participants’ reactions to congruent stimuli, in baseline, SD, and recovery is actually lower than to incongruent ones. It is therefore plausible that, due to a conflict in the incongruent trials, participants took longer to make a response, which at the same time led them to being more accurate. Considering the sleep condition factor, the overall response times to stimuli showed a decreasing trend with the progression of the experiment, that is were the longest during baseline, then SD, and the shortest in recovery, which might be a result of learning. At the same time, participants made slightly more errors when identifying the congruent, as opposed to incongruent stimuli, in each sleep condition, however the differences were small and failed to reach statistical significance. Sleep deprivation and the following recovery period did not influence the number of errors.

The most important finding seems to relate to participants’ omission rates, which differed significantly between the sleep conditions, such that the amount of missed responses increased significantly during the sleep deprivation period, which is in line with the literature that suggests lapses in attention and thus more omissions as a result of sleep deprivation (Lamond et al., 2007; McCarthy & Waters, 1997; van Dongen et al., 2003). Interestingly, the number of omissions decreases significantly in the recovery period, and remains similar to the one seen at baseline.

### Neural effects

There was a significant difference in the participants’ neurophysiological response between the three different sleep conditions of the experiment, such that the amplitude of the P300 differed significantly between baseline and sleep deprivation, and baseline and recovery periods. Interestingly, we did not find a difference between sleep deprivation and recovery, which suggests that after a prolonged period of sleep restriction, one week of a typical sleep routine might not be enough to fully return to the pre-experimental functioning. On the other hand, the P300 amplitude decreases, or in other words habituates, after repeated presentation of the stimulus, as the task becomes somewhat automatic and thus attentional processes might be reduced (Ravden & Polich, 1998, 1999). However, Oginska et al. (2014) have shown that the P300 amplitude reduces after one week period of sleep deprivation. In their study, EEG measurements were taken at four different times a day, after a week of either partial sleep deprivation or unrestricted sleep. Participants were randomly assigned to one of the two groups, one which began the experiment from the sleep deprivation period, followed by a period of unrestricted sleep, and one that had the reverse order. They have found a substantial P300 amplitude reduction following a period of sleep restriction, irrespective of the experimental group, that it regardless of whether the participants first took part in the sleep deprivation or in the unrestricted sleep condition. Those results suggest that the reduction of the P300 amplitude is a result of sleep reduction, rather than a simple matter of habituation.

Despite the very clear negative consequences linked with sleep loss, it has been suggested that the human brain can to some extent adapt to a ‘new normal’ following a period of partial sleep loss. Lamond and colleagues (2007) investigated the processes of recovery with respect to the number of sleep hours (either 9 or 6 or unrestricted recovery following 1 or 2 nights of total sleep deprivation) in the 5-day recovery period. The results showed that only one night of a 9-hour sleep was enough for the subjects’ response speed, lapses, and feeling of sleepiness to return back to the baseline. However, crucially, even though after 2 nights of total sleep loss, and a recovery of 6 hours per night, participants’ performance and feeling of sleepiness failed to return to the baseline, it also did not show an ongoing decline, instead stabilizing at a lower-than-initial but, nonetheless, steady levels. This suggests that reduced but steady sleep recovery conditions after sleep deprivation allow for the human brain to undergo flexible changes. Further evidence suggesting that the human brain has the ability to adapt to different levels of functioning following sleep loss comes from Belenky and colleagues (2003), who did the opposite manipulation to the study described above. Specifically, rather than a variation in the recovery phase, they manipulated the duration of sleep for 7 days (with subjects sleeping either 3, 5, 7, or 9 hours each night), and then looked at the recovery processes (3 nights of 8-hour sleeps). They have found that reduced sleep was linked with some decline in their functioning (such as longer RTs and more lapses of attention). Subjects that in the recovery period slept for 5 or 7 hours per night showed decreased but stable performance in the SD period, whereas those with the biggest sleep curtailment in the SD condition showed perpetually lower performance throughout the whole investigation. Furthermore, it was found that in the group that slept the least in the SD period, both RTs and lapses already improved after the first night of recovery, but from then onwards were comparable to those seen in participants that slept for 5 or 7 hours (that is, who showed steady levels of RTs and lapses in the recovery period, similar to those observed in the sleep deprivation part). Therefore, those subjects showed no enhancement in their behaviour over what was observed during the 7-day period of sleep restriction. Only individuals that had 9 hours of sleep each night of the recovery period showed no differences in either SD and recovery periods, compared to their baseline. Those results further suggest that the human brain can adjust to extended period of sleep curtailment, resulting in performance that, although undoubtedly poorer than initially, is good enough to allow fairly stable functioning.

The strengths and limitations of our study need to be addressed. First, to the best of our knowledge, this is the first EEG study, that looked at chronic sleep restriction over a prolonged period of time, including a week-long recovery phase. There seems to be a gap within the field of partial sleep deprivation research that we have aimed to partially fill. Therefore, despite the clear weaknesses of this study, we believe that it does indeed address a very real societal problem, from both behavioural and neural perspectives. The study suffers from all the weaknesses of a ‘natural experiment’ – there is a number of uncontrollable factors that might affect participants’ lifestyle, alertness, and mood. Furthermore, as it was revealed by the actigraphy, not all the subjects were able to comply strictly with their reduced sleep schedules – there were a few single nights when they overslept. Nevertheless, we decided to keep their data in the analyses. Furthermore, due to feasibility reasons, we did not include a control group, so individuals who would take part in the study but without any external sleep manipulations. Our study sample is also relatively small for an EEG study. On the other hand, we believe that due to the nature of study and the extreme demands it placed on the participants and experimenters alike, it is a reasonable group size. Finally, as it is known from past research (Rupp et al., 2012; van Dongen & Belenky, 2009) that there are robust inter-individual differences in the vulnerability to sleep deprivation. Stable, trait-like inter-individual differences are observed both in responses to total sleep deprivation and in repeated exposures to chronic sleep restriction, in cognitive performance, sleepiness, fatigue (Dennis et al., 2017). Those individual factors were not taken into account in our study and it should be the matter of further explorations.

## General conclusions

The results show that some consequences of chronic partial sleep deprivation can be observed with the use of a Stroop task in a longitudinal study of sleep curtailment and recovery. Our results emphasize that the processes underlying the recovery following an extended period of sleep restriction are more complex and extend far beyond a simple act of increasing the amount of sleep. Furthermore, the long-term consequences of partial sleep deprivation might have been undervalued and should be emphasized both clinically and on a societal level. In the 21st century, a vast proportion of a developed world suffers on a regular basis from chronic partial sleep deprivation. Our results are relevant in terms of further understanding its consequences. There is clearly a need for further studies in this area, particularly the ones that focus on the possible mechanisms involved in the recovery from sleep loss.

## Acknowledgements

This work was supported in part by National Science Center (ncn.gov.pl) grants no., *2011/01/B/HS6/00446*, DEC-2015/17/D/ST2/03492 (JKO); Polish Ministry of Science and Higher Education 7150/E-338/M/2015 and 7150/E-338/M/2016 (KO).

## Supplementary Materials

**Figure 5.**
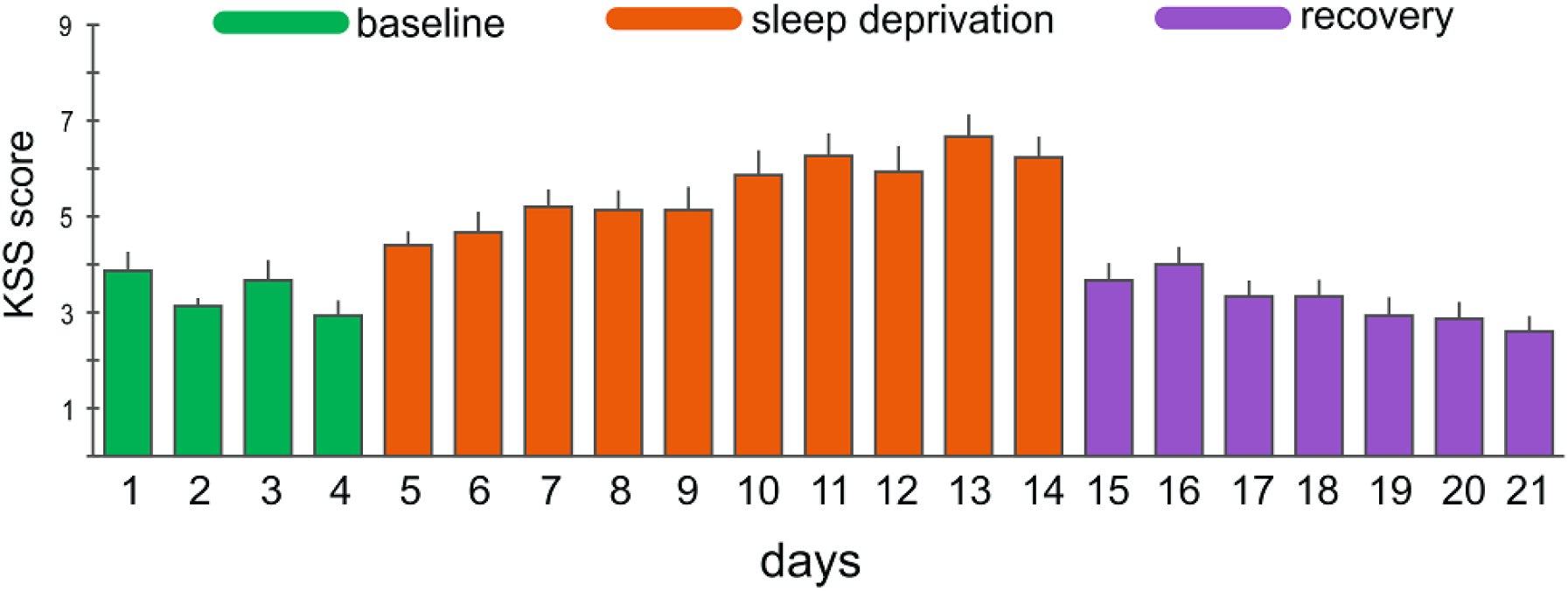
Participants’ KSS scores across the 21 days of the study (n = 14).

**Figure 6.**
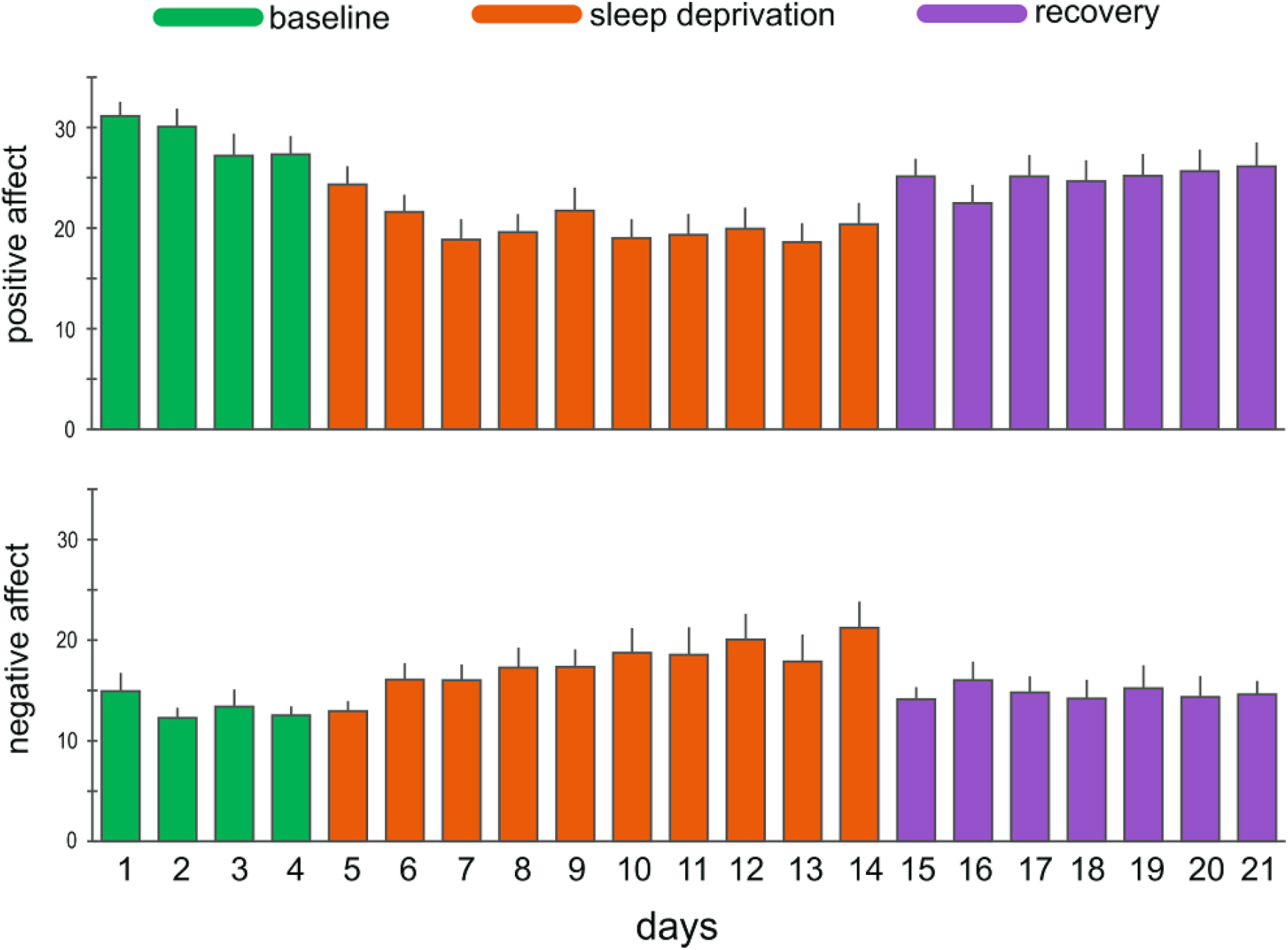
Participants’ PANAS scores across the 21 days of the study (n = 14).

**Figure 7.**
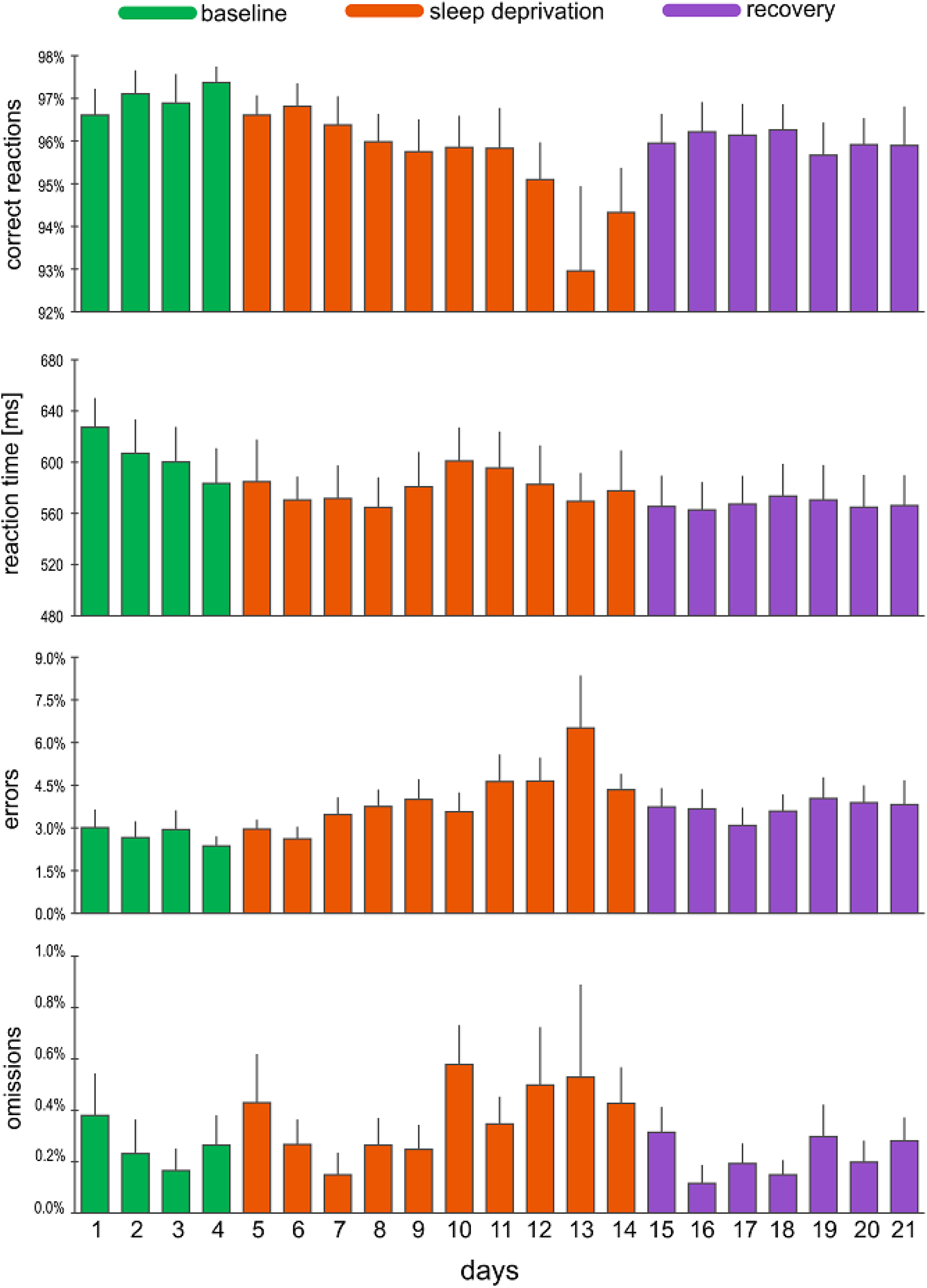
Participants’ behavioural performance across the 21 days of the study (n = 14).

